# Reconstructing the demographic history of blacklegged ticks (*Ixodes scapularis*) in the northern United States

**DOI:** 10.64898/2026.03.10.710853

**Authors:** Dahn-young Dong, Sean D. Schoville

**Affiliations:** Department of Integrative Biology, University of Wisconsin-Madison, Madison, Wisconsin, USA; Department of Entomology, University of Wisconsin-Madison, Madison, Wisconsin, USA

**Keywords:** Approximate Bayesian Computation, genetic demography, historic gene flow, range expansion, vector, blacklegged ticks

## Abstract

**Aim:** To resolve the topological branching patterns, the timing of demographic events, and the effective population size changes associated with major demographic events.

**Location:** Midwestern (eastern North Central) and Northeastern USA

**Taxon:** Blacklegged tick, *Ixodes scapularis* (Say, 1821)

**Methods:** Using three independent genomic datasets, single-nucleotide variants were analyzed for demographic inference. Maximum likelihood topologies and prior ecological knowledge were used to generate nested demographic hypotheses. The best-fit scenario and the associated demographic parameter estimates were determined using approximate Bayesian computation under a random forest statistical model. The topologies and parameters supported in the three independent datasets were compared to generate insights about the demographic history of blacklegged ticks in the region.

**Results:** The emergence of extant northern populations of blacklegged ticks began between 10-15 k.y.a. (thousand years ago), with independent population splits from the common ancestor during the Early-Mid-Holocene, and never more recent than 4 k.y.a. All populations sustained moderately large population sizes without bottlenecks, with Michigan as the exception. Michigan appears to have an uncertain placement that depends on sampling, reflecting its admixed origin.

**Main conclusions:** There are multiple populations of northern blacklegged ticks that have persisted independently as deglaciated regions in the northern U.S. were recolonized following the Last Glacial Maximum (26.5 to 19 k.y.a.). The current ecological expansions across the northern U.S. are likely seeded by separate relictual populations with distinctive genomic ancestry rather than a range expansion from a single source, with important implications for vector-borne disease management.

## Introduction

Efforts to improve our understanding of pest demography can lead to better outcomes in pest management, including more effective monitoring, mitigation, and intervention strategies. However, many pest species are experiencing population expansion events that are difficult to reconstruct from ecological data, as they occur rapidly with sparse monitoring and may involve complex spatiotemporal dynamics that occurrence data may not explain. Genetic data can provide unique historical insights into these demographic events, allowing inferences about the timing of expansion, the source of expanding populations, and the population size changes that accompany spatial range expansion. Several recent examples have successfully leveraged genomic-scale data to reconstruct the complex history of invasive species and endemic pests undergoing spatial expansion events (Barker et al., 2017; Chapuis et al., 2020; Cohen et al., 2022; Eyer et al., 2021; Lombaert et al., 2014; Ryan et al., 2019; Sherpa et al., 2024).

Many species of the hard tick family (*Ixodidae*) are non-permanent parasites that specialize in feeding on vertebrates, disperse via hosts (Beati & Klompen, 2019), and are geographically widespread (Guglielmone et al., 2014). Some have recently undergone population expansion events (Beati et al., 2012) and become pest-like. For example, the rapid growth and spread of the Castor bean tick (*Ixodes ricinus)* in Europe and the blacklegged tick (*I. scapularis* Say, 1821) in the USA, along with a surge in vector borne disease incidence, raises public health concerns (Centers for Disease Control and Prevention, 2024; Eisen & Eisen, 2023; Prevention & Control, 2025; Røed et al., 2016).

Blacklegged ticks, the focus of this study, are native species currently found in the eastern US and central and eastern Canada (Canada, 2022; Centers for Disease Control and Prevention, 2024). Within its range, the northern tick is recognized as a capable vector of the Lyme disease-causing bacterium (*Borrelia burgdorferi sensu stricto*) (Ginsberg et al., 2021). On the other hand, the southern ticks, although the same species, are less abundant and are not currently important vectors for Lyme disease (Ginsberg et al., 2021; Xu et al., 2020). The northern ticks have rapidly increased in number and spread geographically in the Northeastern US since the 1950s, in the Midwestern USA since the 1970s (Figure 1), and more recently in Canada (Clow et al., 2017; Eisen & Eisen, 2023).

**Figure 1.**
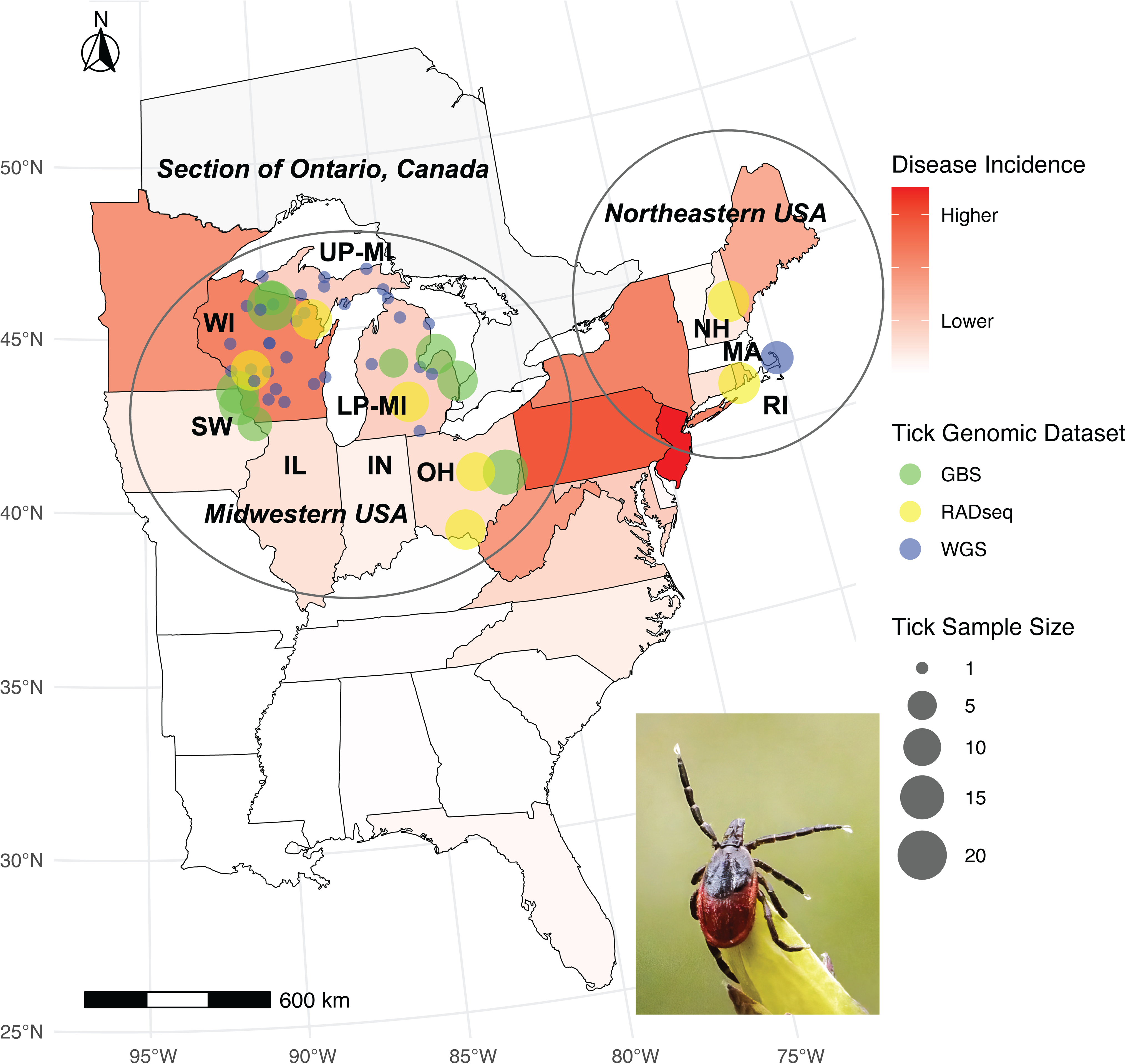
A map of select U.S. states where the blacklegged ticks (*Ixodes scapularis*), the main vector for Lyme disease spirochetes, are collected. The northern USA is the region of the country where Lyme disease is the most common (cases per 100K people in 2021). Wisconsin (WI), Southwest Wisconsin (SW), the Upper and Lower Peninsulas of Michigan (UP/LP-MI), Illinois (IL), Indiana (IN), and Ohio (OH) are part of the Midwestern region, with diverse blacklegged tick genetic ancestry (Dong et al., 2025). Massachusetts (MA), New Hampshire (NH), and Rhode Island (RI) contain tick populations that represent the northeastern genetic cluster (Frederick et al., 2023). The colored dots represent the sampling sites that constitute the populations named after states, while the colors indicate independent genomic datasets. The exception is the three green-highlighted sampling sites adjacent to the SW label, which constitute the SW population. The incidence count data comes from the Centers for Disease Control and Prevention’s WONDER database (Centers for Disease Control Prevention, 2021). Conus Albers Equal Area (EPSG:5070) is used for projection. The photograph of the female adult *I. scapularis*, embedded in the lower right of the figure, is courtesy of Graham Hickling and used with permission.

Long before the recent range expansion, blacklegged ticks may have been present in some northern regions. The northern ticks diverged from the southern ticks at least 50,000 years ago (k.y.a.) (Schoville et al., 2024). During the Last Glacial Maximum (LGM) from 26.5 to 19 k.y.a. (Clark et al., 2009), when glaciers covered parts of the northern U.S. (Dalton et al., 2022), the northern populations may have been restricted to the Atlantic coastal refugia (Whitmore et al., 1967; Xu et al., 2020), experiencing significant demographic contraction (Schoville et al., 2024). It is possible that Northeastern populations later migrated after the last glacier’s retreat to form the Midwestern populations and had shared historical gene flow (Xu et al., 2020), even though current gene flow between these regions is limited (Humphrey et al., 2010). After the LGM, northern populations likely rebounded but may have fluctuated due to extensive deforestation and declines in host populations, such as deer, which reached their smallest size around 130 years ago, leaving probable small relictual tick populations that went unnoticed in the northern region (Eisen & Eisen, 2023). Over the past decades, the reemergence of the northern blacklegged tick has appeared invasive: much of the northern U.S. was largely free of ticks 100 years ago, and now ticks are almost ubiquitous in the region (Centers for Disease Control and Prevention, 2024) (Figure 1).

The current extent of Midwestern and Northeastern populations remains an active area of research regarding whether the recent population increase, range expansion, and associated rise in Lyme disease incidence resulted from local reemergence of relictual tick populations at low population densities or from effective dispersal of a few key source populations. In the Northeast, it is believed that current northward dispersal from a common source population is the main mode of range expansion, rather than reemergence of multiple relictual populations (Khatchikian et al., 2015). In contrast, in the Midwest, genetic evidence suggests that range expansions came from multiple isolated sources, as gene flow diminishes at geographical scales of ∼300 kilometers (Dong et al., 2025). Specifically, Wisconsin may not be the source of other Midwestern states, contrary to what has been proposed (Eisen & Eisen, 2023). A new, distinct population in southwestern Wisconsin has been identified, and populations in Michigan, a neighboring state, are largely genetically distinct from other Midwestern populations (Dong et al., 2025), prompting a reexamination of the source of the active range expansion events. In addition, parsing out the influence of historical admixture and current gene flow, both of which influence patterns of neutral variation in the genome, could advance our understanding of the forces shaping tick biology across different time scales (He et al., 2013). In particular, the importance of dispersal and local adaptation in tick population dynamics (Frederick et al., 2023; Schoville et al., 2024) would inform whether tick management should focus on distinct "management units" (Dong et al., 2025).

Integrating three recent population genomic datasets, and with the temporal and genomic flexibility of approximate Bayesian computation (ABC) for demographic analyses (Beichman et al., 2018), we ask: what is the sequence of branching events that give rise to the current northern populations, what is the timing of events, and were population size changes involved during the process of divergence and expansion?

## Methods

### Genetic Sampling

Previously published genomic data from the Midwest include a genotype-by-sequencing (GBS) dataset (NCBI BioProject PRJNA1193100, (Dong et al., 2025)), a whole-genome sequencing (WGS) dataset (NCBI BioProject PRJNA731189, (Schoville et al., 2024)), and a RADseq (restriction site-associated DNA sequencing) dataset (NCBI BioProject PRJNA852262 (Frederick et al., 2023)). Blacklegged ticks (nymphs and adults) in the GBS and WGS datasets were sampled while questing (host-seeking) via dragging (Falco & Fish, 1992) from 2021 to 2023 across the Midwestern United States. The blacklegged ticks in the RADseq dataset were collected from a mixture of dragging and animal hosts from 2010 to 2021.

From the GBS dataset (Dong et al., 2025), four genetically distinct populations were selected based on geographical relevance and sample size: one from the cluster in Wisconsin (WI_GBS_), one representing the cluster in southwestern Wisconsin, Iowa, and Indiana (SW_GBS_), one representing Michigan (MI_GBS_), and one representing a genetically distinct population in Ohio (OH_GBS_). Specifically, in WI_GBS_, we combined the nearby Kemp Natural Resource Research Station (n=20) and North Trout Lake (n=8) samples in north-central Wisconsin as a single population (n=28), as they have a low FST (0.005). For the SW sample, we combined Dubuque (n=8), Lansing (n=14), and McGregor (n=12) to represent a single population (n=34), as they are geographically close and genetically similar (FST < 0.008). In MI_GBS_, we combined samples from Lakeport State Park Campground (n=12), CMU Neithercut Woodland (n=5), and Sleeper State Park (n=12), all located in central-eastern Michigan, into a single population (n=29; FST < 0.009). Lastly, in OH_GBS_, we selected Tappan Lake Park (n=16) from southeastern Ohio as the representative population (Figure 1).

Our goal is to infer demographic patterns from the GBS results and to leverage WGS and RADseq as replicate datasets for comparison. To compare the GBS results with the WGS dataset, we obtained samples from Wisconsin (WI_WGS_, n=19), Michigan (MI_WGS_, n=14), and the Northeastern US, represented by Massachusetts (NE_WGS_, n=7) (Figure 1). These represent geographically separated blacklegged tick samples from the northern US (Schoville et al., 2024). The WI_WGS_ samples may correspond to two population clusters (WI_GBS_ and SW_GBS_) in the GBS dataset because the sample size in the SW_GBS_ region was too small to analyze separately. Our purpose is to estimate divergence among regions outside Wisconsin over time. By combining samples, we assume WI_WGS_ populations follow a structured coalescent model, in which the divergence time to NE_WGS_ is not expected to be influenced by combining closely related WI_WGS_ population clusters (Nielsen & Wakeley, 2001; Strasburg & Rieseberg, 2010).

Similarly, from the RADseq dataset, we focus on samples from Wisconsin (WI_RAD_, n=24, from Marinette County population and Monroe County population with FST = 0.01), one Michigan site (MI_RAD_, n=12, from Ingham County), Ohio (OH_RAD_, n=23, from Knox County population and Scioto County population with FST = 0.02), and a Northeastern US site (NE_RAD_, n=26, combined NH and RI, FST = 0) (Figure 1).

### Genomic data processing for the three genomic datasets

Raw GBS data were processed and filtered using a previously described workflow using a reference genome (NCBI: GCF_016920785.2) (Dong et al., 2025), with minor modifications to filter missing data. Briefly, variant calling was conducted following the best practices outlined in the Germline Short Variant Discovery pipeline, utilizing GATK 4.4.0.0 (Van der Auwera & O’Connor, 2020). All variants were restricted and reformatted as unphased biallelic single-nucleotide polymorphisms (SNPs) to apply missingness filters. Various variant-missingness thresholds (<0.20) were applied using VCFTOOLS 0.1.17 (Danecek et al., 2021) to retain high-quality variants. The resulting SNPs were filtered by depth of coverage (DP), retaining an average DP of 15-100 to improve variant quality. Imputation was performed with BEAGLE 5.4 (Browning et al., 2018) to predict missing variants using linkage information. The imputed dataset was then subjected to linkage pruning (window size 50kb, step size 10, R-squared threshold 0.2) with PLINK2 v2.00a3 SSE4.2 (Chang et al., 2015) to exclude variants that could bias demographic analysis.

The raw WGS data and RADseq data were processed similarly with some modifications. WGS data were originally generated with moderate coverage (∼5x), so while they were similarly processed using GATK and filtering was used to remove variants with missing data, DP filtering was set between 5 and 100, and the genome was subset to putatively neutral variants, using SNPEFF and SNPSIFT to select intergenic SNPs (Cingolani, Patel, et al., 2012; Cingolani, Platts, et al., 2012). Additionally, linkage pruning was performed in R using ‘snprelate’ v 1.34.1 (max base pairs in sliding window: 2000, max SNPs in sliding window: 50, linkage disequilibrium threshold = 0.2) (Zheng et al., 2012), instead of PLINK2, because the total sample size was fewer than 50. Raw RADseq data were also processed similarly to the GBS pipeline, except that variants with missing genotypes were removed, and linkage pruning was performed in R using ‘snprelate’ instead of PLINK2 because the sample size in some subsets was fewer than 50.

To conduct nested demographic analyses for the genomic data, three Variant Call Format (VCF) files were separately ascertained from the GBS dataset with an increasing number of populations: 1. first SNP set called from OH_GBS_ and SW_GBS_; 2. second SNP set called from OH_GBS_, SW_GBS_, and WI_GBS_; and finally, 3. third SNP set was called from OH_GBS_, SW_GBS_, WI_GBS_, and MI_GBS_. In a similar fashion, the WGS dataset was used to generate two SNP sets, one including NE_WGS_ and WI_WGS_, and one including NE_WGS_, WI_WGS_, and MI_WGS_. By the same token, the RADseq dataset generated three SNP sets: one including NE_RAD_ and WI_RAD_; one including NE_RAD_, W_RAD_, and OH_RAD_; and one including NE_RAD_, WI_RAD_, OH_RAD_, and MI_RAD_.

### Demographic hypotheses generation

Without prior knowledge, a large number of demographic hypotheses can be proposed about the evolutionary relationships among populations, including many possible topologies (branching orders), timing, changes in effective population size, and potential admixture events. However, testing many hypotheses through extensive simulations is impractical. Therefore, we initially limited hypothesis generation by using TREEMIX 1.13 (Pickrell & Pritchard, 2012), a method that infers maximum-likelihood population topologies and gene flow events. We converted VCF files with more than two populations from the three genomic datasets into TREEMIX inputs using STACKS 2.64 (Catchen et al., 2013) and generated the most likely tree(s) for each dataset, each run with 100 replicates. We inferred potential admixture events for the SNP sets, allowing 0 to 3 migration edges. If any number of migration edges was indicated as a better fit than no migration, we used the three-population (Reich et al., 2009) and four-population TREENESS test (Keinan et al., 2007) in TREEMIX to verify whether the migration edge was statistically significant. Specifically, in the TREENESS test, a significantly negative *f*3 statistic suggests one admixed population, whereas a significantly non-zero *f*4 value suggests gene flow. The best-supported tree hypotheses were then used to design demographic models that could be tested with approximate Bayesian computation (ABC) inference. To gain a more comprehensive understanding of demographic relationships, we compared the TREEMIX demographic hypotheses with several other ecological hypotheses generated *a priori* based on previous knowledge of the blacklegged ticks in the sampled region (Eisen & Eisen, 2023; Lantos et al., 2017).

### Approximate Bayesian statistics for demographic inference

#### Nested inference

ABC uses population genetic summary statistics to compare simulated and observed data across various demographic models, with the aim of identifying the best-fitting model and estimating parameters of interest (Beichman et al., 2018). Past research on the range expansion of different species has employed a nested approach to demographic modeling (Estoup & Guillemaud, 2010; Rovito & Schoville, 2017; Ryan et al., 2019). This nested analysis builds iteratively on the best-fitting initial models by adding populations while preserving the prior ranges of the existing parameters, gradually increasing the number of free parameters to create a more robust test of demographic history. Specifically, we begin by estimating the most supported demographic model for two populations that are likely the most divergent among all populations. Then, a more recently derived population is included, and the evolutionary relationship of this new population is tested with the existing ones until all populations have been evaluated with optimized relationships. One important assumption we made about the two population models is that the two populations emerged independently from a common ancestor rather than directly from each other, as constrained by TREEMIX (Figure S 1).

We began with GBS-Analysis 1, conducting two-population demographic inference between OH_GBS_ and SW_GBS_, which are the most divergent based on the TREEMIX analysis (Figure S 1d) and contain more genetic diversity (Dong et al., 2025). Analysis 2 performs a three-population demographic inference including WI_GBS_. In Analysis 3, a four-population demographic inference is conducted by including MI_GBS_. The parameter values were estimated only for the best-supported scenario across the full set of populations. Similarly, WGS and RADseq analyses also began with TREEMIX analyses to generate *a priori* topologies and migration hypotheses, constrained by nested ABC tests with a plausible set of hypotheses that allow for an increasing number of populations organized by the presumed order of divergences.

#### Pipeline construction

Among various ABC implementations, DIYABC-Random Forest (DIYABC-RF) v1.0 was chosen for its flexible visualization, command-line access, and intuitive model comparison framework (Collin et al., 2021). Due to specific SNP input file requirements, the sample order in each VCF file was first reorganized by population using BCFTOOLS 1.16 (Danecek et al., 2021), and then converted into DIYABC-RF-compatible “.snp” files with VCF2DIYABC (Loire, 2015). Because of the large number of SNPs and strict filtering steps, we expect our data to be effectively neutral, so no additional filters were applied at this stage. We set sex assignment as “not specified,” marked as "9." We used a cutoff of 15000 SNPs for the GBS dataset, 20000 SNPs for the RADseq and WGS datasets, as the software manual states that analyzing between 5000 and 20000 SNP loci typically yields robust results. Similarly, the manual indicates that, because the SNP dataset is biallelic, the mutation model is simplified so that only one mutation could have occurred in the coalescence tree of sampled genes, allowing the default Hudson’s simulation algorithm to speed up the simulation (Collin et al., 2021). This allows us to avoid specifying a mutation rate for *I. scapularis*. To convert time units from generations to years, we set the generation time of the blacklegged tick to 2 years based on the typical life cycle (Keirans et al., 1996).

All three genomic datasets were used to independently test demographic scenarios, generate training datasets, perform model choice, and estimate demographic parameters. Most of these steps are user-defined, except for the summary statistics, which are automatically selected based on the number of populations tested. The summary metrics are: the proportion of monomorphic loci, heterozygosity for individual populations and pairs, FST for populations, pairs, triplets, quadruplets, and overall population, as well as Peterson’s f3-statistics, f4-statistics, Nei’s distance for each pair of populations, and the maximum likelihood coefficient of admixture for each triplet of populations (Table S 1). Model specifications are detailed in the input header files, created using the graphical user interface version of the DIYABC-RF.

#### Prior assignments

The model parameters and their ranges of values need to be specified *a priori* to simulate data under the hypothesized topology (Table S 2). We began with the same parameter ranges for all genomic datasets, but the ranges of the shared prior parameters across WGS and RADseq datasets vary slightly from those of GBS due to fine-tuning during initial simulations to ensure compatibility with the observed data.

Nevertheless, the priors overlap, ensuring comparability of all parameters across the three genomic datasets. At the same time, during nested inference within each dataset, the parameters that have already been fine-tuned remain the same for the more complex model analyses.

For example, for the GBS dataset, the split times of existing populations were constrained between 5 and 10000 generations ago. This broad range allows the best-fit scenario to optimize possible population split times, ranging from recent range expansions (within a few hundred years to the present) (Eisen & Eisen, 2023) to historical divergence (more than a few hundred years ago) (Schoville et al., 2024). The effective population size was set between 1000 and 1 million, reflecting a wide range.

Note that the effective population size is an idealized measure that assumes a Hardy-Weinberg equilibrium and represents the same level of genetic drift as the actual population, which differs from the census size. We chose this large range because, although some blacklegged tick populations are undergoing rapid range expansions with potential bottlenecks, many established populations likely exist in substantial numbers given their high reproductive rate and diverse host preferences (Oliver Jr et al., 1993). Short-term bottlenecks were not explicitly tested because negative Tajima’s D values (Dong et al., 2025) suggested expanding population sizes, indicating no significant bottlenecks. Instead, we modelled changes in population size during major population splits to describe population dynamics.

#### Initial and final inferences

An initial assessment of prior distributions and model compatibility with the observed data was performed by simulating 500 datasets per scenario and analyzing the percentage of overlap between the observed and simulated data’s summary statistics. When a sufficient proportion of simulated data scenarios exhibit significant overlap with the observed data across the majority of summary statistics (>5%), the hypothesized scenario and the prior parameters are deemed compatible. Next, each compatible scenario was simulated with 20,000 datasets for model choice and with 110,000 simulations for parameter estimation. These numbers of simulations are at the upper bound for sufficiency to obtain robust results, according to the DIYABC user manual.

Random forest parameter estimation uses 100,000 simulated datasets with 1500 classification votes to determine the best parameter values, with accuracy metrics computed from 10,000 out-of-bag datasets, which are considered optimized.

We focus on estimating parameters from the best-scenario that includes the largest number of populations from each of the three datasets, i.e., GBS-Analysis Three (four populations), WGS-Analysis Two (three populations), and RADseq-Analysis Three (four populations). For each parameter, the median, 90% quantile range, and local (posterior) median NMAEs (normalized mean absolute error) are calculated. Local NMAEs are indicators of the relative interpretability of model parameters (Chapuis et al., 2020; Collin et al., 2021), with values below one considered interpretable. The lower the local NMAE, the greater our confidence in the interpretation of the parameter value. All model choice analyses and parameter estimations are averaged over ten replicates to ensure stability and account for variation. The final demographic models and associated parameter estimates were visualized using DEMES (Gower et al., 2022).

## Results

### Genomic data processing and demographic hypotheses testing

For three genomic datasets, multiple VCF input files were created. The assessment of missingness, mean variant depth, and number of SNPs per file is listed in Table S 3.

The number of SNPs generated for each VCF file exceeds 10000, with varying SNP counts, given variant quality and genomic diversity of the datasets. We retained more WGS SNPs than the other two datasets to be more inclusive of diverse alleles and to reflect greater genomic divergence among populations.

To limit the number of tested demographic scenarios, we used TREEMIX to generate 100 maximum-likelihood trees for the three- and four-population SNP sets for each genomic dataset, with inferred optimized migration events, if any (Figure S 1). The GBS three-population SNP set revealed three equally likely topologies without migration (Table S 4) (topology 1 is excluded for ABC testing due to nested inference constraint), while the four-population SNP set revealed a plausible topology with a migration event (66% replicate support) but was not statistically significant in the TREENESS test. The WGS three-population SNP set also revealed three equally likely topologies without migration. Similarly, the RADseq three-population SNP set revealed three plausible topologies without migration, and the four-population SNP set revealed one plausible topology with one statistically significant migration event, supported by the TREENESS test, which has two alternatives (each with 54% and 44% support across replicates). Both possible migration edges were further tested in ABC. The TREENESS migration significance test is detailed in (Table S 5).

The scenarios adopted from TREEMIX, along with other ecologically plausible scenarios tested using ABC, are displayed in Table 1 and visualized in Figure S 2. For GBS-Analysis 2 (GBS-A2), in addition to three scenarios compatible with the TREEMIX (Scenarios 1-3), Scenario four was created as a modification of Scenario two, assuming that WI_GBS_ and OH_GBS_ arose around the same time from the common ancestor. For GBS-A3, in addition to models consistent with TREEMIX (Scenarios one and two), we created Scenario three to position MI_GBS_ as the closest sister population to WI_GBS_. This alternative scenario is based on ecological research that MI has recently been populated by blacklegged ticks, possibly from a WI source (Lantos et al., 2017). For WGS-A2, in addition to three scenarios compatible with TREEMIX (Scenarios 1-4), we created Scenario five that positions MI_WGS_ as a possible product of admixture between WI_WGS_ and NE_WGS_. RADseq-A2 is constructed similarly to WGS-A2. For RADseq-A3, all scenarios are compatible with the TREEMIX without additions, as possible admixture is already accounted for by TREEMIX.

**Table 1.** Model choice results of nested demographic inference of the three genomic datasets (GBS, WGS, and RADseq) for the blacklegged tick populations using approximate Bayesian computation – random forest implementation (ABC-RF). All results are reported as averages with standard deviations in parentheses after ten replicates. For each analysis, the best scenario and its results are shown in bold type. The best scenario was subsequently used to build the next nested analysis within each genomic dataset. Across replicates, all analyses consistently voted for the same best scenarios, except for GBS-Analysis 1, with 80% support, and WGS-Analysis 1 and RAD-Analysis 2, with 90% support. Posterior probability is reported for the winning scenario of each analysis. Note that the same population acronyms used across genomic datasets do not indicate the same samples nor the exact population. See the method section of the main text and Figure 1 for details. Visual representation of the compared scenarios is shown in Figure S 2.

| Nested Analysis | Prior error % (SD) | Scenario ID and description | RF votes (SD) | Posterior % (SD) |
| --- | --- | --- | --- | --- |
| GBS-A1 | 0.26 (0.003) | S1: OH arose from the CA first | 644 (89) | - |
|  |  | **S2: SW arose from the CA first** | **856 (89)** | **0.55 (0.039)** |
| GBS-A2 | 0.26 (0.001) | S1: WI arose from the CA before other populations | 127 (24) | - |
|  |  | S2: WI arose from the CA before OH and after SW | 222 (37) | - |
|  |  | **S3: WI arose from SW** | **818 (60)** | **0.72 (0.051)** |
|  |  | S4: WI and OH split the same time from the CA | 333 (38) | - |
| GBS-A3 | 0.091 (0.001) | **S1: MI is the most closely related to OH** | **804 (39)** | **0.72 (0.034)** |
|  |  | S2: MI is the most closely related to OH, with admixture from the MI ancestor to SW | 583 (34) | - |
|  |  | S3: MI is the most closely related to WI | 114 (12) | - |
| WGS-A1 | 0.31 (0.003) | S1: NE arose from the CA first | 663 (70) | - |
|  |  | **S2: WI arose from the CA first** | **837 (70)** | **0.54 (0.047)** |
| WGS-A2 | 0.23 (0.001) | S1: MI arose from the CA first | 43.4 (21) | - |
|  |  | S2: MI arose from the CA between WI and NE | 45.4 (9.3) | - |
|  |  | **S3: MI arose from WI** | **1120 (51)** | **0.74 (0.018)** |
|  |  | S4: MI arose from NE | 3.90 (2.4) | - |
|  |  | S5: MI arose from WI and NE admixture | 285 (49) | - |
| RAD-A1 | 0.39 (0.003) | **S1: NE arose from the CA first** | **914 (53)** | **0.60 (0.043)** |
|  |  | S2: WI arose from the CA first | 587 (53) | - |
| RAD-A2 | 0.18 (0.0007) | S1: OH arose from the CA first | 534 (48) | - |
|  |  | S2: OH arose from the CA between WI and NE | 148 (20) | - |
|  |  | **S3: OH arose from NE** | **684 (60)** | **0.83 (0.026)** |
|  |  | S4: OH arose from WI | 13 (5.1) | - |
|  |  | S5: OH arose from WI and NE admixture | 122 (21) | - |
| RAD-A3 | 0.40 (0.002) | S1: MI arose after its ancestor's split from WI and admixture with OH | 142 (15) | - |
|  |  | S2: MI arose after its ancestor's split from WI and admixture with NE | 487 (37) | - |
|  |  | S3: MI arose after its ancestor's split from the CA and admixture with OH | 195 (19) | - |
|  |  | **S4: MI arose after its ancestor's split from the CA and admixture with NE** | **676 (51)** | **0.60 (0.023)** |

Notes: RAD: RADseq; SD: standard deviation; RF: random forest; CA: common ancestor; NE: Northeastern USA, WI: Wisconsin; SW: Southwest Wisconsin; MI: Michigan; OH: Ohio

Table 1 shows the model choice for each analysis across three genomic datasets. Averaged results from ten replicates show that the prior error rate for each analysis set ranged from 0.09 to 0.40, with the posterior probability of the best-fit model ranging from 0.54 to 0.83. The first analysis (A1) across genomic datasets typically shows a lower posterior probability in the winning scenario, whereas later nested analyses show a higher posterior probability. From the best-fit model for each genomic dataset, it is clear that multiple independent population emergences occurred from the common ancestor of these northern blacklegged ticks Figure S 2. However, there are some topological differences: Michigan’s origin shows varying support across three genomic datasets, with possible admixture as a potential explanatory factor.

### Inferred parameter estimates

Based on the best supported, most complex scenario for each genomic dataset, GBS-Analysis 3, WGS-Analysis 2, and RADseq-Analysis 3, respectively (Table 1), we obtained estimates for the models’ demographic parameters (Figure 2, Table S 6). The common ancestor of the sampled extant populations experienced an approximately 60-fold expansion in population size around 70-160 k.y.a (the effective population size expanded from 3-7k to 160-420k). The emergence of distinct northern populations of blacklegged ticks was supported by all three models and began between 10-15 k.y.a. (thousand years ago), with additional population splits from the putative common ancestor throughout the Early-Mid-Holocene period, though never more recent than 4 k.y.a. All extant populations sustained moderately large population sizes since their common ancestor (27k to 570k in Table S 6) with Michigan clearly experiencing a bottleneck.

**Figure 2.**
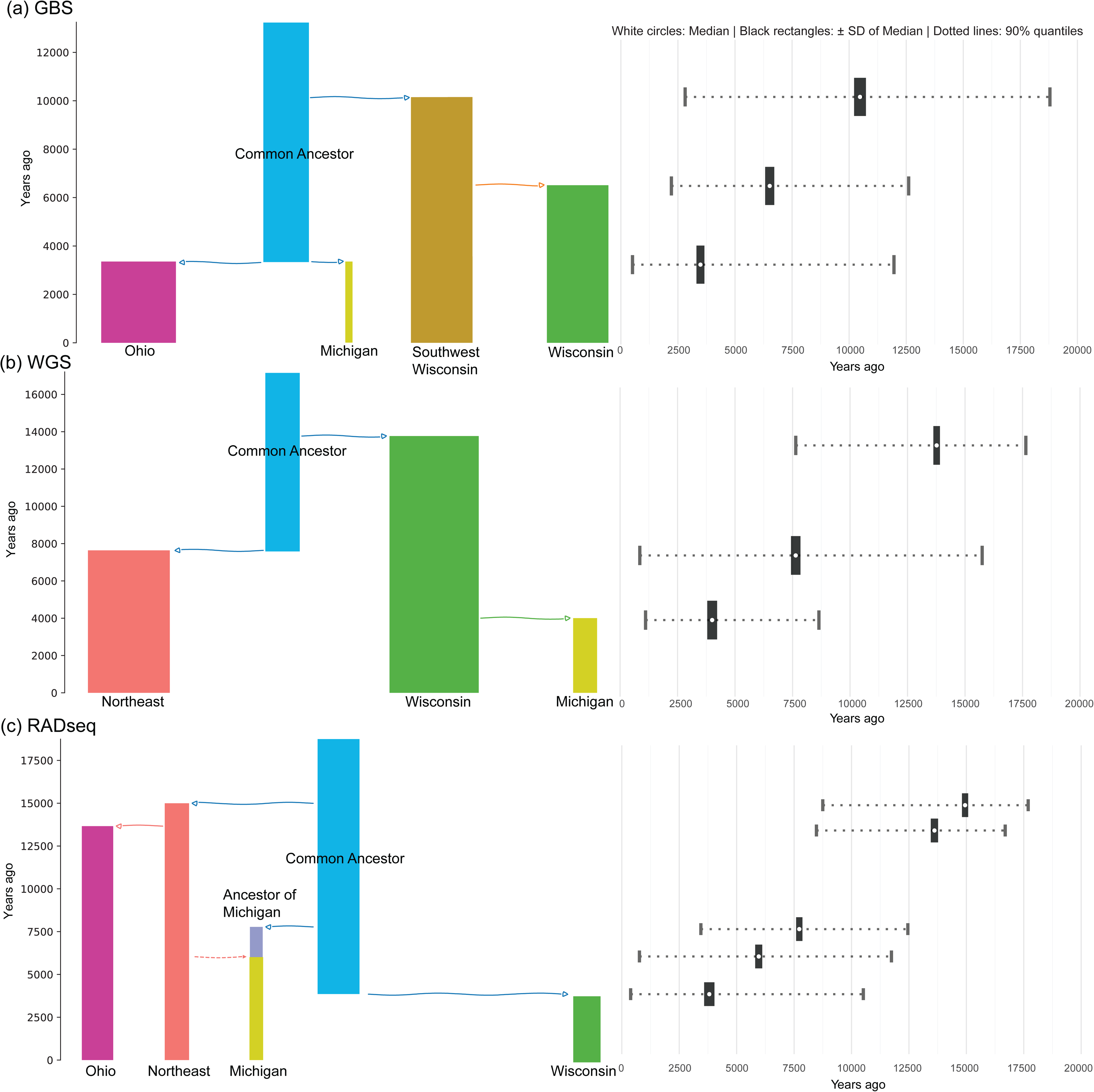
Topologies and demographic parameter estimates for blacklegged tick populations under the best-supported demographic scenario, based on three independent genomic datasets, with values averaged across ten replications. Panels (a) to (c) correspond to results from GBS, WGS, and RADseq genomic datasets, respectively. The left column indicates the best demographic history for each dataset, and the right column shows the detailed parameter estimates corresponding to the major demographic events, aligned with the arrows in the left column. Solid arrows represent population splits, while color changes indicate changes in population identity. A dashed arrow indicates a pulse of admixture event. Arrow length does not carry meaning. The widths of the population bars in the left column indicate the relative effective population sizes for the populations within each panel. Note that the same acronyms used across genomic datasets do not indicate the same samples nor the exact population. See the method section of the main text and Figure 1 for details.

The Midwestern populations (Wisconsin, Michigan, and Ohio) exhibit various relationships among themselves across the genomic datasets, although Michigan consistently emerged the latest (Figure 2). Furthermore, Michigan’s relationship with Wisconsin, its close neighbor, is challenged by its frequent disassociation from Wisconsin and affiliation with populations closer to the northeastern US (Northeast and Ohio). Based on these best-supported scenarios across three genomic datasets, a few discrepancies remain. The relative timing of emergence of the Wisconsin and Northeast populations switches positions (in the GBS and WGS datasets, Wisconsin emerged early, whereas in the RADseq dataset, it emerged late). Similarly, Michigan’s origin is ambiguous: it appears more distant from the Wisconsin population in GBS and RADseq datasets, whereas Wisconsin is shown as the origin for Michigan in the WGS dataset.

Nevertheless, if we contextualize the extant population split time results across all three genomic datasets (Figure 2) by focusing on the parameters’ local NMAEs with the highest support (Table S 6), results suggest that Wisconsin emerged from the common ancestor of northern population 14 k.y.a. (WGS dataset with NMAE = 0.24), Michigan emerged from Wisconsin 4 k.y.a. (WGS dataset with NMAE = 0.46), Ohio emerged from the Northeast 14 k.y.a (RADseq dataset with NMAE = 0.16), and the Northeast emerged from the common ancestor of northern population 15 k.y.a (RADseq dataset with NMAE = 0.18).

## Discussion

We set out to reconstruct the demographic history of the northern blacklegged tick populations in the U.S., leveraging recent GBS data (Dong et al., 2025), as well as WGS and RADseq data (Frederick et al., 2023; Schoville et al., 2024), which allow for the reconstruction of relationships and split times of the tick populations in the Midwestern and Northeastern regions. In particular, we tested hypotheses regarding the branching patterns, the timing of demographic events, and the population size changes associated with major demographic events. The improved understanding of the demographic history among these populations clarifies the historical dispersal events over time and elucidates how the spread of blacklegged ticks may have occurred.

Improved knowledge may change how we perceive the rapid range expansion of this native parasite in the region, and what we can do to control its populations and the spread of tickborne disease.

Our results suggest that the earliest rise of blacklegged ticks in the Midwest region did not originate directly from the Northeastern population, unlike what has been hypothesized in the literature (Humphrey et al., 2010). Furthermore, the northern populations likely have multiple origins from a common ancestor rather than dispersal from a single source, as previously suggested (Khatchikian et al., 2015). In terms of the timing of events, most extant populations derived from a common ancestor and have since expanded around 10-15 k.y.a. (Figure 2). This indicates that tick populations have persisted independently and undergone separate range expansions as they recolonized deglaciated regions in the north. The ecological expansions currently being observed across the northern U.S. are likely seeded by separate relictual populations with distinctive genomic ancestry. Furthermore, genomic differences dating to Early-to-Mid-Holocene divergences, rather than recent gene flow across regions, are best supported by the data and indicate that population structure is still being preserved.

### Full model specifics and interpretations

#### Branching order of extant populations: multiple sources and Northeastern connection

The subdivision within Wisconsin was investigated with GBS-ABC analysis (Figure 2a). The WI_GBS_ emerged from SW_GBS_, thereby situating the spatial origin of the recent Midwestern reemergence farther south (Figure 1). While it is still likely that some relictual population existed in northern Wisconsin and contributed to a Wisconsin-wide expansion in recent decades (Eisen & Eisen, 2023), it is clear that the WI_GBS_ is not the region’s oldest lineage, but rather that the SW_GBS_ is the oldest.

The source of the Michigan region receives varied support, consistent with previous findings that Michigan lacks genetic clustering with the other midwestern populations (Dong et al., 2025). ABC modeling, more often than not, places Michigan populations at a topological location farther from the quintessential Midwest populations and closer to either Ohio, which carries non-Midwest genetic signals, or to Northeastern populations via ancient admixture (Eisen et al., 2016; Frederick et al., 2023). The variation among datasets might be best explained by admixture, which can lead to gene-tree population-tree discordances (Peter, 2016). Our result continues to support the idea that there are multiple sources of relictual populations that persisted over time and re-emerged in recent conditions (Hoen et al., 2009), and that Michigan populations have been distinct from other extant populations for thousands of years.

Lastly, blacklegged ticks are increasingly being found in Ohio and are believed to be admixed populations as Midwestern ticks expanded eastward and Northeastern ticks expanded westward (Frederick et al., 2023). The Ohio samples we tested, however, are collected from the eastern half of the state and are more likely related to northeastern populations (Figure 2c).

#### Timing of emergence of extant populations: Early-Mid-Holocene origins rather than contemporary expansion

We selected broad prior ranges for the split times of the sampled populations (Table S 2) to encompass the possibility of recent range expansion, as public health data suggested (Eisen & Eisen, 2023), or ancient population divergences (capped at the time of the LGM) despite the recent reemergence. The results are insightful because none of the split-time estimates suggest a recent range expansion between regions; rather, the genetic signals of a range shift extend as far back as 15 k.y.a., from the end of the Pleistocene to the early Holocene (Figure 2). This indicates that the northern populations represent local relictual populations that have undergone separate recent expansion events. Patterns within states often reflect recent spatial expansion with contemporary gene flow (Dong et al., 2025), but across the northern U.S., ancestral population genetic structure persists.

It is possible that the Holocene Climatic Optimum (9-6 k.y.a.) created suitable habitats (Chapuis et al., 2020) for the expansion of the common ancestor across the northern US, leading to lineage splitting for some populations, as glaciers had fully retreated from the region (Dyke, 2004). The split times are also consistent with the existing literature, which places any northern population subdivisions within the past 50K years (Schoville et al., 2024) and recent population growth in the Wisconsin region within 20 k.y.a. (Van Zee et al., 2015).

#### Population size changes: stable expansions

Most extant populations are large, with limited evidence of founding events and bottlenecks, as seen in classic invasion biology. Most size changes between and after population-split events are not drastic (Figure 2, Table S 6). The result that populations are large and stable, except for Michigan, suggests that most Early-Mid-Holocene divergences were gradual expansions that led to large contemporary populations. The Michigan population originated from a small initial size from the common ancestor, which is still evident today in its genetic homogeneity (Dong et al., 2025). It is important to note that some confidence intervals for a few parameters are unreliable (Table S 6: NMAE > 1). This might be due to data limitations or model misspecification. However, such an outcome is not unusual in the literature (Berdugo-Cely et al., 2023; Eyer et al., 2021). It is reassuring, though, that across three genomic datasets, the timing of major splitting events and population sizes had high support, and all estimates fall within similar geological ranges (Table S 6).

#### Timing of the common ancestor size change

The common ancestor that gave rise to the sampled extant populations expanded 4-fold in effective size at approximately 70 to 150 k.y.a. (Table S 6), which occurs close to the last interglacial period. A similar population expansion event was identified during this period through whole-genome sequence analysis using sequential Markov coalescent methods (Schoville et al., 2024). It should also be noted that Schoville et al. (2024) estimated the north-south divergence at 50 k.y.a., which suggests that our estimates of population expansion pertain to an ancestor that includes southern *I. scapularis* populations. This result is also consistent with the Bayesian extended skyline analysis of populations in North Carolina, often categorized as the front line of the north-south division, which expanded 200 k.y.a. (Van Zee et al., 2015). Future studies should not only investigate the genetic subdivisions of the northern populations, especially the location and timing of refugia for the northern populations (Xu et al., 2020), but also the geographical and temporal sequence of the “north-south” divergence.

### Historic and current gene flow in distinguishing between the emergence of native populations and the spread of invasive species

In determining patterns of recent gene flow among blacklegged tick populations in the Midwest, previous research has shown effective intra-regional dispersal, but limited inter-regional dispersal (Dong et al., 2025). In line with this result, genetic demographic inference of the same population clusters revealed ancient range shifts dating back to the Early-Mid-Holocene that created the larger-scale population genetic structure, with separate lineages evident over the past 4000 years (Figure 2). It is important to acknowledge that the genetic composition of current populations is the product of both past and present gene flow, and that this gene flow can be detected and described from robust spatial sampling (Chapuis et al., 2020; He et al., 2013). From the perspective of the blacklegged tick’s natural history in the U.S., it is becoming clear that Midwestern ticks are likely to have recently re-emerged from multiple relictual populations that originated thousands of years ago and have gone undetected, rather than being invaded by a single recent source. In addition, the current literature presents different estimates of recent gene flow between the Midwestern and Northeastern ticks (Humphrey et al., 2010; Van Zee et al., 2015). Our result suggests that Midwestern ticks are related to Northeastern populations, but only through shared ancestry rather than through recent gene flow.

Our results shed new light on the biogeographical boundary between the Northeastern and Midwestern ticks, as Michigan’s non-Midwest identity is supported by the majority of the genomic dataset, and Ohio’s early split from the Northeast is evident. The biogeographic patterns of other widespread species in the same region in North America has suggested ring-like routes of range expansion and patterns of admixture around the Great Lakes, where Ohio populations may maintain genetic connectivity with the Northeast via a Northern Appalachian refugium and Michigan population via southern Ontario, Canada (Kim et al., 2018; Lee-Yaw et al., 2008; Waldron et al., 2024). Future work should sample key transition areas, such as southern Ontario and the vicinity of the Appalachian Mountains, to better understand the historical and contemporary influences of Northeastern ticks, and to more explicitly parse out the genetic influence of geological and climatic events from contemporary events.

The study was initially motivated by ascertaining the sequential and numerical timing of the Midwestern blacklegged tick range expansion events that appeared invasive: much of the northern U.S. was largely free of ticks one hundred years ago and now ticks are almost ubiquitous in the region (Centers for Disease Control and Prevention, 2024), with Lyme disease spreading broadly and increasing rapidly (Figure 1). However, our estimates of divergence between sampled populations suggest that split times are much more ancient, and relictual populations in the Midwestern region remained unnoticed (Dong et al., 2025; Hoen et al., 2009; Humphrey et al., 2010). While most invasive species experience founder effects (bottlenecks with significant genetic drift), blacklegged tick populations have not. The populations have remained genetically diverse, likely have considerable adaptive capacity, and may exhibit important local differences (Schoville et al., 2024). These results show the importance of reconstructing demographic history, even in well-studied species, as assumptions about ecological dynamics might be overly simplified (Cohen et al., 2022; Ortego et al., 2021), and it is not always clear what constitutes a local versus an invasive species (Sherpa et al., 2024).

### Implications for blacklegged ticks and Lyme disease management

The insight that Midwestern blacklegged tick populations have multiple genetic origins dating back to the Early-Mid-Holocene time can inform us about current population dynamics. Our research suggests that relictual populations have recently repopulated the region, perhaps as secondary forest developed and the white-tailed deer (*Odocoileus virginianus*) population rebounded (Adams & Hamilton, 2011). With the uptick of Lyme disease incidence and knowledge that blacklegged ticks are the main vector for this disease in the region, we can infer that these independently arising relictual populations, with their unique mix of moderately high genetic diversity, all retain the ability to sustain and transmit the pathogen among hosts. It is likely that, when blacklegged ticks were constrained in relictual populations, they still circulated pathogens among wildlife hosts such as rodents (Prado et al., 2022), but did not do so extensively across geographic space. Although blacklegged ticks are rapidly expanding in numbers and geographic range, they still maintain genetic breaks across geography. This isolation across populations may suggest that each population, with distinct ancestry, harbors unique phenotypes adapted to local environmental conditions (Schoville et al., 2024). Thus, tick populations at larger geographical scales might be treated as distinct management units that require population-specific monitoring of the evolution of pesticide resistance, as these populations have been evolving separately for a few thousand generations. At the same time, the Lyme disease causative agent, *Borrelia burgdorferi,* is more genetically homogenous than blacklegged ticks (Humphrey et al., 2010), one of its suitable vectors. Therefore, leveraging the pathogen’s genetic homogeneity may be a more effective option for Lyme disease control than targeting the blacklegged ticks.

## Data Availability Statement

Scripts, genotype data, and input data that were used to generate analysis results are available on Dryad (DOI link available after peer review)

## Acknowledgements

We thank Dr. Paskewitz, Dr. Xia Lee, and other members of the Paskewitz Lab for providing most of the Wisconsin GBS samples. We thank Michelle Volk and Dr. Jean Tsao from Michigan State University for collecting Michigan GBS samples, as well as the Michigan Department of Natural Resources, Central Michigan University, Ann Arbor Parks and Recreation, and St. Joseph County Parks and Recreation. We thank Green Lee from the Indiana Department of Health and Lee’s assistants for collecting GBS samples from Indiana, and Allison Williams and Dr. Risa Pesapane from the Parasite and Pathogen Ecology (PPE) Lab at Ohio State University for collecting GBS samples from Ohio. We also thank Dr. Ryan Smith and his colleagues at Iowa State University for providing a list of potential sites to visit in Iowa to collect GBS samples. We thank Michael Troutman, Zach Farrand, Ebony Dominique Taylor, and Emma Terris for their written feedback. A generative AI–based coding assistant (Anthropic’s Claude Sonnet, implemented in GitHub Copilot) was used to help draft and debug portions of the R/Python codebase and bash scripts, including utility functions for extracting outputs across replicate runs, data wrangling, and computing standard summary statistics. All processing steps and analytical decisions were defined by the authors, and all code was manually reviewed, tested, and validated; the authors take full responsibility for the results. Funding was provided by a USDA McIntire-Stennis award (#WIS04035).

