## Supplemental File for "Reconstructing the demographic history of blacklegged ticks (*Ixodes scapularis*) in the northern United States"

### Dryad data accessibility reviewer link:

### <http://datadryad.org/share/LINK_NOT_FOR_PUBLICATION/If9onGcqwJkDzTWZQN-h_QmxkWJmpFpOEF2kIWHZdzw>

[Figure S 1. Panel a-c shows the three most likely three-population topologies inferred from the GBS dataset using TREEMIX maximum-likelihood analysis, based on 100 replicates. Panel d displays the four-population topology of the GBS dataset, with an ancestor of the Michigan population admixed with the Southern Wisconsin population. Similarly, panel e-g shows the three-population topologies inferred via TREEMIX replicates for the WGS dataset. Panels h-j show the three-population topologies for the RADseq dataset, and panels k and l show the four-population topologies from the RADseq dataset, with an ancestor of the Michigan population admixed with either the Ohio or the Northeast population.	4](#_Toc223947167)

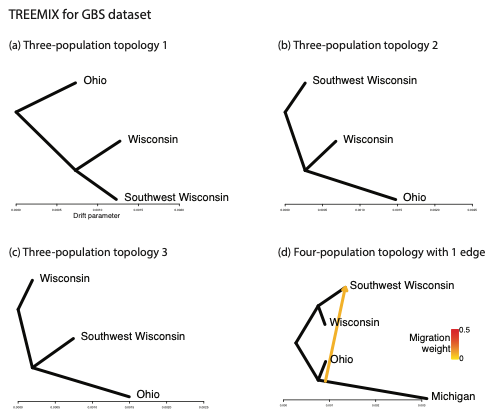

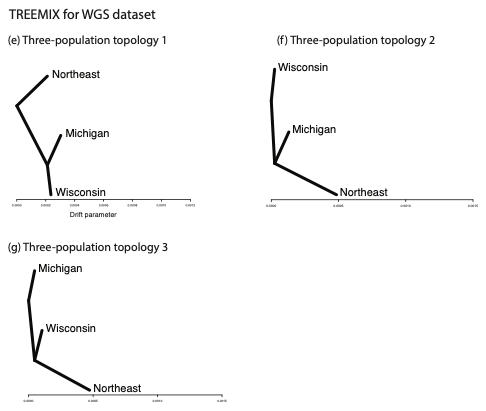

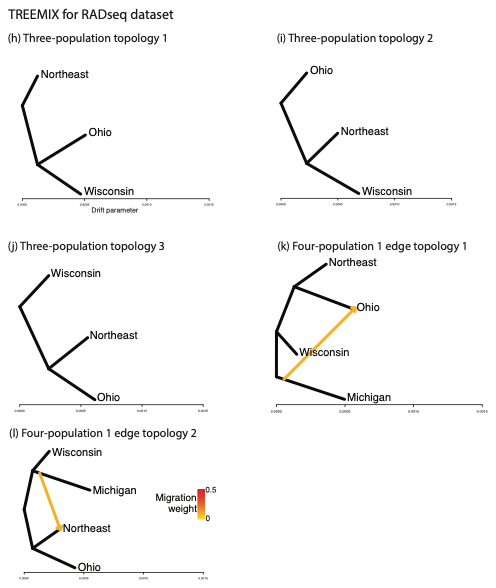

Figure S 1. Panel a-c shows the three most likely three-population topologies inferred from the GBS dataset using TREEMIX maximum-likelihood analysis, based on 100 replicates. Panel d displays the four-population topology of the GBS dataset, with an ancestor of the Michigan population admixed with the Southern Wisconsin population. Similarly, panel e-g shows the three-population topologies inferred via TREEMIX replicates for the WGS dataset. Panels h-j show the three-population topologies for the RADseq dataset, and panels k and l show the four-population topologies from the RADseq dataset, with an ancestor of the Michigan population admixed with either the Ohio or the Northeast population.

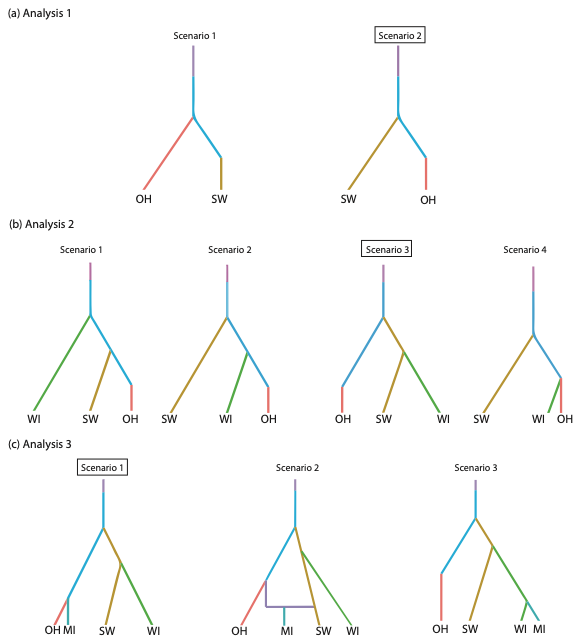

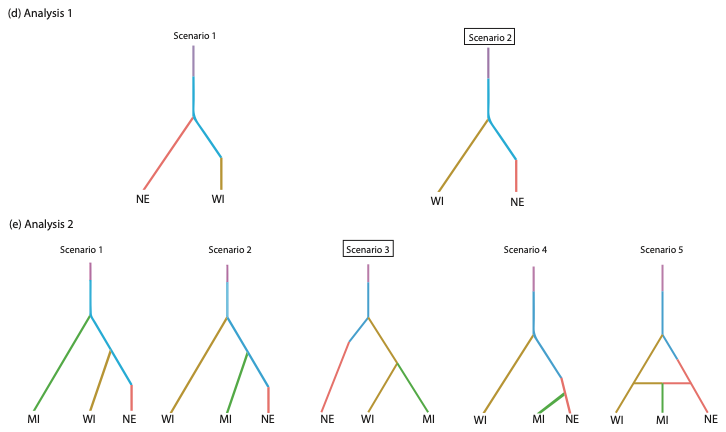

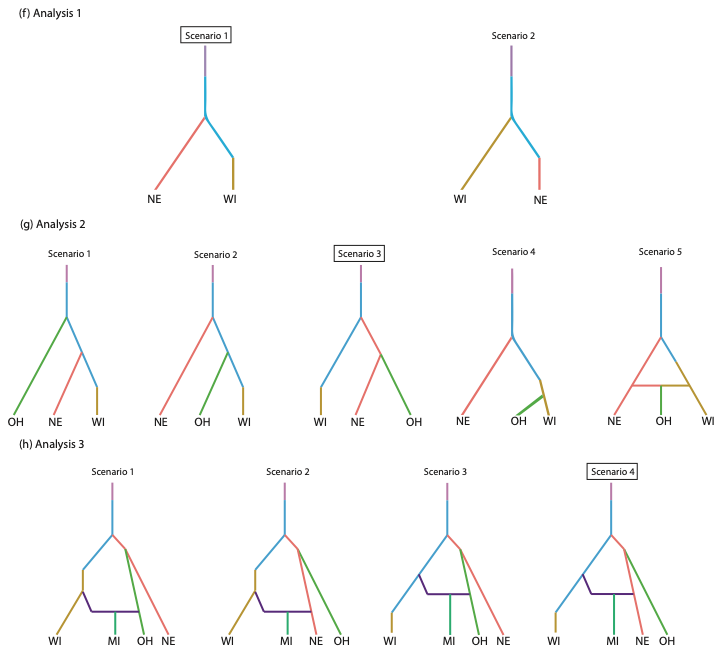

Figure S 2. Coalescent scenarios modeled for ABC-RF analysis. Panels a-c, d-e, and f-h are nested analyses for GBS, WGS, and RADseq genomic datasets, respectively. Black bounding boxes indicate the winning scenario for each analysis. Each color represents a modeled population size. Horizontal edges indicate admixture events. The sampled extant populations are labeled with population acronyms. Note that the same acronyms used across genomic datasets do not indicate the same samples nor the exact population. See the method section of the main text and Figure 1 for details.

Table S 1. Summary statistics tested based on the number of sampled populations tested.

| Two-population (16 statistics) | Three-population (50 statistics) | Four-population (130 statistics) |
| --- | --- | --- |
| ML1p 1 2  HWm 1 2  HWv 1 2  HBm 1.2  HBv 1.2  FST1m 1 2  FST1v 1 2  FST2m 1.2  FST2v 1.2  NEIm 1.2  NEIv 1.2 | ML1p 1 2 3  ML2p 1.2 1.3 2.3  HWm 1 2 3  HWv 1 2 3  HBm 1.2 1.3 2.3  HBv 1.2 1.3 2.3  FST1m 1 2 3  FST1v 1 2 3  FST2m 1.2 1.3 2.3  FST2v 1.2 1.3 2.3  NEIm 1.2 1.3 2.3  NEIv 1.2 1.3 2.3  AMLm 1.2.3 2.1.3 3.1.2  AMLv 1.2.3 2.1.3 3.1.2  FST3m 1.2.3  FST3v 1.2.3  F3m 1.2.3 2.1.3 3.1.2  F3v 1.2.3 2.1.3 3.1.2 | ML1p 1 2 3 4  ML2p 1.2 1.3 1.4 2.3 2.4 3.4  ML3p 1.2.3 1.2.4 1.3.4 2.3.4  HWm 1 2 3 4  HWv 1 2 3 4  HBm 1.2 1.3 1.4 2.3 2.4 3.4  HBv 1.2 1.3 1.4 2.3 2.4 3.4  FST1m 1 2 3 4  FST1v 1 2 3 4  FST2m 1.2 1.3 1.4 2.3 2.4 3.4  FST2v 1.2 1.3 1.4 2.3 2.4 3.4  NEIm 1.2 1.3 1.4 2.3 2.4 3.4  NEIv 1.2 1.3 1.4 2.3 2.4 3.4  AMLm 1.2.3 2.1.3 3.1.2 1.2.4 2.1.4 4.1.2 1.3.4 3.1.4 4.1.3 2.3.4 3.2.4 4.2.3  AMLv 1.2.3 2.1.3 3.1.2 1.2.4 2.1.4 4.1.2 1.3.4 3.1.4 4.1.3 2.3.4 3.2.4 4.2.3  FST3m 1.2.3 1.2.4 1.3.4 2.3.4  FST3v 1.2.3 1.2.4 1.3.4 2.3.4  FST4m 1.2.3.4  FST4v 1.2.3.4  F3m 1.2.3 2.1.3 3.1.2 1.2.4 2.1.4 4.1.2 1.3.4 3.1.4 4.1.3 2.3.4 3.2.4 4.2.3  F3v 1.2.3 2.1.3 3.1.2 1.2.4 2.1.4 4.1.2 1.3.4 3.1.4 4.1.3 2.3.4 3.2.4 4.2.3  F4m 1.2.3.4 1.3.2.4 1.4.2.3  F4v 1.2.3.4 1.3.2.4 1.4.2.3 |
| Full names of the acronyms. Copied directly from DIYABC-RF 1.0 user manual  “1. [ML1p] [ML2p] [ML3p] [ML4p] - Proportion of monomorphic loci for each population, as well as for each pair, triplet and quadruplet of populations.  Mean (m suffix added to the code name) and variance (v suffix) over loci values are computed for all subsequent summary statistics  2. [HWm] [HWv] [HBm] [HBv] - Heterozygosity for each population and for each pair of populations…  3. [FST1m] [FST1v] [FST2m] [FST2v] [FST3m] [FST3v] [FST4m] [FST4v] [FSTGm] [FSTGv] - FST-related statistics for each population (i.e., population-specific FST; Weir & Goudet 2017), for each pair, triplet, quadruplet and overall populations (when the dataset includes more than four populations) …  4. [F3m] [F3v] [F4m] [F4v] – allele shared Patterson’s f-statistics for each triplet (f3-statistics) and quadruplet (f4-statistics) of populations…  5. [NEIm] [NEIv] - Nei’s (1972) distance for each pair of populations  6. [AMLm] [AMLv] - Maximum likelihood coefficient of admixture computed for each triplet of populations...” | | |

Table S 2. Prior parameters and summary statistics tested by DIYABC-RF across analyses for three genomic datasets

| **GBS ABC parameters** | **Distribution** | **Prior parameter range** |
| --- | --- | --- |
| Emergence timing of the sampled populations, and timing of admixture (generations ago) | Uniform | 5-10000 |
| Emergence timing of the most recent common ancestor population (generations ago) | Uniform | 10000-100000 |
| Population sizes of Ancestors (Effective Population size) | Log-uniform | 1000-1000000 |
| Population size of sampled population (Effective Population size) | Log-uniform | 1000-1000000 |
| Population size of admixed population (Effective Population size) | Uniform | 1000-1000000 |
| Rate of admixture | Uniform | 0.05-0.2 |

| **WGS ABC parameters** | **Distribution** | **Prior parameter range** |
| --- | --- | --- |
| Emergence timing of the sampled populations (NE and WI) (generations ago) | Uniform | 50-9000 |
| Emergence timing of the most recent common ancestor (generations ago) | Uniform | 9000-90000 |
| Population size of Ancestor (Effective Population size) | Log-uniform | 1000-1000000 |
| Emergence timing of the sampled population (MI) and timing of admixture | Uniform | 100-15000 |
| Population sizes of the most recent common ancestor (Effective Population size) | Log-uniform | 45000-800000 |
| Population size of NE (Effective Population size) | Log-uniform | 200000-1500000 |
| Population size of WI (Effective Population size) | Log-uniform | 100000-750000 |
| Population size of MI (Effective Population size) | Log-uniform | 20000-500000 |
| Rate of admixture | Uniform | 0.01-0.99 |

| **RADseq ABC parameters** | **Distribution** | **Prior parameter range** |
| --- | --- | --- |
| Emergence timing of the sampled populations (NE, WI, and MI) and timing of admixture (generations ago) | Uniform | 50-9000 |
| Emergence timing of the most recent common ancestor (generations ago) | Uniform | 9000-90000 |
| Population size of Ancestor (Effective Population size) | Log-uniform | 1000-1000000 |
| Emergence timing of the sampled population (OH) | Uniform | 10-9000 |
| Population sizes of the most recent common ancestor (Effective Population size) | Log-uniform | 45000-800000 |
| Population size of NE and WI (Effective Population size) | Log-uniform | 45000-700000 |
| Population size of OH (Effective Population size) | Log-uniform | 45000-1500000 |
| Population size of MI (Effective Population size) | Log-uniform | 1000-700000 |
| Rate of admixture | Uniform | 0.05-0.5 |

Table S 3. Statistics of the processed genomic data

| Genomic source – Analysis number | Populations involved (population size is listed only once for each population) | Variant missingness allowed (%) | Mean variant depth | SNP retained |
| --- | --- | --- | --- | --- |
| GBS-A1 | OH_GBS_ (n=16) and SW_GBS_ (n=34) | 4 | 31 | 28439 |
| GBS-A2 | OH_GBS_, SW_GBS_, and WI_GBS_ (n=28) | 6 | 33 | 18367 |
| GBS-A3 | OH_GBS_, SW_GBS_, WI_GBS_, and MI_GBS_ (n=29) | 15 | 33 | 11994 |
| WGS-A1 | NE_WGS_ (n=7) and WI_WGS_ (n=19) | 0 | 9 | 215000 |
| WGS-A2 | NE_WGS_, WI_WGS_, and MI_WGS_ (n=14) | 0 | 8 | 311000 |
| RADseq-A1 | NE_RAD_ (n=26) and WI_RAD_ (n=24) | 0 | 34 | 37305 |
| RADseq-A2 | NE_RAD_, W_RAD_I, and OH_RAD_ (n=23) | 0 | 36 | 38919 |
| RADseq-A3 | NE_RAD_, WI_RAD_, OH_RAD_, and MI_RAD_ (n=12) | 0 | 30 | 38360 |

Table S 4. TREEMIX replicate support for three-population topologies. The numbered topologies are not the same across genomic datasets.

|  | Topology 1 | Topology 2 | Topology 3 |
| --- | --- | --- | --- |
| GBS | 31% (OH;WI,SW) | 29% (SW;WI,OH) | 40% (WI;SW,OH) |
| WGS | 35% (NE;WI,MI) | 37% (WI;NE,MI) | 28% (MI;NE,WI) |
| RADseq | 42% (NE;OH,WI) | 36% (OH;NE,WI) | 22% (WI;NE,OH) |

Table S 5. Three and four-population “treeness” significance test for all three genomic datasets. A significantly negative *f*3 statistic suggests one admixed population, whereas a significantly non-zero *f*4 value suggests gene flow.

|  | Population tree topology | F statistics | Standard error | Z score | P value (two sided for 4-pop test and one-sided for 3-pop test) | interpretation |
| --- | --- | --- | --- | --- | --- | --- |
| GBS four-pop | OH,MI;MS,WI | -2.14303e-05 | 5.47092e-05 | -0.391712 | 0.70 | No significant gene flow |
| GBS three-pop | MS;OH,WI | 0.000555623 | 7.53275e-05 | 7.37611 | < .00001 | No significant gene flow |
|  | OH;MS,WI | 0.00148386 | 0.000114384 | 12.9726 | < .00001 | No significant gene flow |
|  | WI;OH,MS | 0.00046893 | 6.22755e-05 | 7.52992 | < .00001 | No significant gene flow |
| WGS three-pop | NE;WI,MI | 0.000434804 | 2.92099e-05 | 14.8855 | < .00001 | No significant gene flow |
|  | MI;NE,WI | 9.63787e-05 | 1.17015e-05 | 8.23641 | < .00001 | No significant gene flow |
|  | WI;NE,MI | 4.73449e-05 | 1.05042e-05 | 4.50723 | < .00001 | No significant gene flow |
| RADseq four-pop | NE,WI;OH,MI | 0.000170685 | 2.95644e-05 | 5.77333 | < .00001 | Significant gene flow |
| RADseq three-pop | NE;WI,OH | 0.000277901 | 2.93647e-05 | 9.46378 | < .00001 | No significant gene flow |
|  | OH;NE,WI | 0.000388868 | 4.47596e-05 | 8.68794 | < .00001 | No significant gene flow |
|  | WI;NE,OH | 0.000384634 | 5.07253e-05 | 7.58267 | < .00001 | No significant gene flow |

Table S 6. Parameter estimates of the best supported scenario for each genomic dataset (GBS, WGS, RADseq). The estimates are presented as the median with the evaluators, the 90% Quantile, and the local NMAE in median form, after ten replications. Note that the same population acronyms (NE: northeast, WI: Wisconsin; SW: Southwest Wisconsin; MI: Michigan; OH: Ohio) used across genomic datasets do not indicate the same samples nor the exact population. See the method section of the main text and Figure 1 for details. Emergence time is indicated by generations ago.

|  | **Averaged Median (SD)** | **Averaged Q05 (SD)** | **Averaged Q95 (SD)** | **Averaged Local Median NMAE (SD)** |
| --- | --- | --- | --- | --- |
| **GBS Parameters** |  | | | |
| EPS of ancient ancestor (NA) | 37300 (2920) | 2250 (481) | 175000 (73900) | 1.72 (0.451) |
| EPS of OH (N1) | 303000 (34100) | 39900 (9100) | 841000 (40400) | 1.6 (0.802) |
| EPS of SW (N2) | 236000 (11200) | 69700 (10400) | 723000 (23100) | 0.492 (0.0648) |
| EPS of WI (N3) | 221000 (7940) | 48400 (3750) | 706000 (33600) | 0.617 (0.149) |
| EPS of MI (N4) | 26700 (1220) | 5130 (738) | 79400 (2270) | 0.701 (0.11) |
| EPS of CA (N5) | 164000 (6610) | 53400 (3250) | 579000 (54200) | 0.446 (0.042) |
| ET of MI and OH (t1) | 1740 (173) | 254 (50.3) | 5980 (287) | 1.45 (0.353) |
| ET of WI (t3_al) | 3260 (200) | 1110 (88.5) | 6300 (294) | 0.449 (0.0882) |
| ET of SW (t2) | 5240 (259) | 1410 (178) | 9400 (80.8) | 0.77 (0.444) |
| ET of CA (ta5) | 77900 (1920) | 34000 (2450) | 98400 (296) | 0.308 (0.0665) |
| **WGS Parameters** | **Averaged Median (SD)** | **Averaged Q05 (SD)** | **Averaged Q95 (SD)** | **Averaged Local Median NMAE (SD)** |
| ET of NE (t1_al) | 3820 (206) | 429 (89.7) | 7870 (125) | 1.53 (0.741) |
| ET of WI (t2_al) | 6880 (143) | 3820 (228) | 8820 (43.7) | 0.239 (0.0294) |
| ET of MI (t3_al2) | 2000 (214) | 554 (95.7) | 4320 (282) | 0.459 (0.0999) |
| ET of CA (ta4) | 36600 (1010) | 18300 (690) | 69600 (3430) | 0.288 (0.0338) |
| EPS of ancient ancestor (NA) | 3040 (188) | 1190 (54.4) | 8390 (746) | 0.315 (0.0242) |
| EPS of NE (N1) | 523000 (36600) | 217000 (4120) | 1380000 (35700) | 0.48 (0.039) |
| EPS of WI (N2) | 570000 (14300) | 284000 (6670) | 736000 (3700) | 0.214 (0.0209) |
| EPS of MI (N3) | 153000 (13800) | 37900 (2620) | 417000 (15400) | 0.371 (0.0293) |
| EPS of CA (N4) | 220000 (26900) | 67500 (5390) | 641000 (36300) | 0.455 (0.065) |
| **RAD Parameters** | **Averaged Median (SD)** | **Averaged Q05 (SD)** | **Averaged Q95 (SD)** | **Averaged Local Median NMAE (SD)** |
| ET of NE (t1) | 7500 (135) | 4400 (214) | 8870 (22.3) | 0.184 (0.0253) |
| ET of WI (t2) | 1930 (224) | 217 (32.4) | 5280 (306) | 1.56 (0.257) |
| ET of OH (t3_al2) | 6830 (160) | 4260 (186) | 8370 (143) | 0.162 (0.0399) |
| ET of ancestor of MI (t4_al) | 3890 (138) | 1750 (90.8) | 6250 (170) | 0.293 (0.0398) |
| ET of CA (ta5) | 51100 (1250) | 30800 (1380) | 75700 (1530) | 0.19 (0.0275) |
| ET of MI and admixture (t6mix) | 3010 (154) | 409 (69.8) | 5890 (202) | 1.24 (0.465) |
| Admixture rate (ra6) | 0.12 (0.01) | 0.05 (0) | 0.33 (0.02) | 0.494 (0.0686) |
| EPS of ancient ancestor (NA) | 7800 (328) | 2640 (402) | 14000 (723) | 0.429 (0.126) |
| EPS of NE (N1) | 250000 (4090) | 154000 (3930) | 358000 (7420) | 0.164 (0.0404) |
| EPS of WI (N2) | 281000 (19000) | 68800 (5870) | 646000 (11600) | 0.608 (0.119) |
| EPS of OH (N3) | 323000 (17400) | 167000 (9980) | 638000 (28800) | 0.181 (0.014) |
| EPS of MI (N4) | 141000 (23000) | 18800 (2140) | 619000 (13900) | 0.919 (0.256) |
| EPS of CA (N5) | 432000 (15800) | 157000 (10900) | 756000 (11400) | 0.269 (0.0203) |
| EPS of the MI ancestor (N6) | 132000 (14200) | 8380 (1460) | 632000 (17900) | 1.71 (0.411) |

Note: CA stands for the common ancestor of all extant populations. EPS stands for effective population size. ET stands for emergence time.
